# The Role of Pavlovian Bias in Smoking Cessation Success and Neurocognitive Alterations During Nicotine Withdrawal

**DOI:** 10.1101/2025.11.18.688678

**Authors:** Eunhwi Lee, Heesun Park, Jeung-Hyun Lee, Hyeonjin Kim, Hyung Jun Park, Hee-Kyung Joh, Woo-Young Ahn

## Abstract

**Background:** Inability to manage conflicts between multiple decision-making systems is associated with various mental disorders including addiction. The imbalance between the Pavlovian and instrumental systems may contribute to smoking cessation failure and relapse, as excessive Pavlovian motivation may override goal-directed quitting attempts. However, the exact role of the Pavlovian bias in smoking cessation and its neurocognitive mechanisms remain poorly understood.

**Methods:** Eighty-six smokers participated in a smoking cessation clinic, completing the well-established orthogonalized go-nogo task before and after the intervention. We applied computational modeling and functional magnetic resonance imaging (fMRI) analyses to investigate the role of Pavlovian bias and its neural correlates in predicting smoking cessation outcomes.

**Results:** Computational modeling revealed that higher levels of Pavlovian bias were associated with treatment failures. Importantly, Pavlovian bias significantly moderated the relationship between clinic participation rate and the likelihood of smoking cessation, suggesting that such bias can undermine motivational effort to quit. Furthermore, individuals who successfully quit smoking exhibited increased Pavlovian bias toward reward following cessation, indicating a potential cognitive mechanism underlying the relapse cycle. Complementary fMRI analyses demonstrated that Pavlovian bias influenced valence encoding in the ventromedial prefrontal cortex (vmPFC), suggesting altered decision-making neural circuits.

**Conclusions:** Pavlovian bias can be a critical risk factor of relapse in the smoking population, while this bias can be increased in response to nicotine abstinence. Moreover, the results highlight the roles of the vmPFC in decision-making under Pavlovian-instrumental conflict, providing insights into the neural underpinnings of relapse vulnerability. Understanding these mechanisms may guide targeted interventions to enhance smoking cessation outcomes.

## Introduction

Decision-making in humans is a complex, value-based process guided by subjective evaluations of available (1,2). Central to this process are two primary learning systems: the Pavlovian system, which is innate and hard-wired, learns stimulus-outcome associations and drives instinctive behaviors such as approaching rewards and avoiding punishments. In contrast, the instrumental system computes stimulus-action-outcome associations, enabling individuals to evaluate the value of specific actions (2,3). These systems typically cooperate, promoting efficient learning and optimal decision-making (4,5). However, conflicts may arise when Pavlovian responses—or ‘biases’—interfere with goal-directed actions governed by the instrumental system, resulting in suboptimal choices and maladaptive behaviors (4,6).

To capture this conflict, previous research has employed paradigms such as Pavlovian-to-Instrumental Transfer (PIT), where reward- or punishment-associated cues influence instrumental actions. Typically, positively conditioned cues enhance approach behaviors, whereas negatively conditioned cues inhibit them (7–9). However, because PIT measures Pavlovian influence only after the learning phase, the orthogonalized go/no-go (GNG) task was developed to assess Pavlovian bias directly during learning by orthogonally separating action (go/no-go) and valence (reward/punishment). Studies using the orthogonalized GNG task have shown that participants perform better in conditions with congruent action-valence pairings (e.g., go to win, no-go to avoid loss), highlighting the role of Pavlovian bias in learning and decision-making (10).

Various studies have investigated the neural correlates of Pavlovian bias and instrumental control using PIT and GNG paradigms, particularly focusing on the role of cortico-basal-ganglia loops. The striatum, a core component of these loops, is central to reward-based learning by signaling reward prediction errors (RPEs) (11–13). This further encodes instrumental action values and guides action selection in Pavlovian and instrumental learning (12,14–16). GNG studies have revealed that striatal and substantia nigra/ventral tegmental area (SN/VTA) track action value signals and encode ‘go’ requirements, suggesting their roles in behavioral invigoration (10,17,18). Meanwhile, cortical regions such as the ventromedial prefrontal cortex (vmPFC) and anterior cingulate cortex (ACC) have been implicated in valence encoding and in inhibiting Pavlovian responses (19–23).

As mentioned earlier, Pavlovian bias may constitute a risk factor for maladaptive behaviors and psychiatric conditions. Of particular interest is substance use disorders (SUDs), which are strongly tied to dysfunctional reward processing and maladaptive alterations in dopaminergic signaling within striatal and midbrain circuits (24–26). These neurocognitive changes are closely linked to Pavlovian motivation, where individuals exhibit compulsive approach tendencies and strong ‘wanting’ of rewards despite suboptimal long-term outcomes, a process heavily influenced by Pavlovian conditioning (4).

Previous studies provide mixed evidence of Pavlovian bias as a significant factor in SUDs. Some studies using the PIT paradigm have shown that patients with alcohol dependence (AD) exhibit greater Pavlovian influence compared to healthy controls and PIT-related BOLD signals predicted prospective alcohol intake and relapse (27–29). On the other hand, some studies have reported that patients with AD showed intact PIT effects (30–32). In a study investigating the PIT among patients with nicotine use disorder (NUD), their nicotine dependency was not associated with Pavlovian bias (33).

We hypothesize that addressing the motivational state or conflict in SUDs might be crucial for understanding the role of Pavlovian bias. As mentioned, Pavlovian bias may lead to suboptimal behaviors mainly when it conflicts with the goal-directed responses led by the instrumental system (4). If an individual lacks motivation to quit, Pavlovian bias may have little effect since there is no conflict between the Pavlovian responses and instrumental goals. However, when motivated to quit, automatic responses to addiction-related cues can hinder the cessation efforts. Thus, considering the individual’s motivational state and its interaction with Pavlovian bias might be essential to fully understand its role in SUDs. Previous studies have often failed to account for these conflicts, which may contribute to the mixed findings. This study aims to address this gap by examining the interactions between Pavlovian bias and cessation attempts in predicting cessation outcomes.

Furthermore, although previous studies have linked dopamine to Pavlovian bias (34,35), no longitudinal study to our knowledge reported altered Pavlovian bias following long-term abstinence. Abstinence is known to induce alterations in neurotransmitter systems, such as decreases in dopamine levels (36). The SN/VTA, which are crucial for action value processing, are closely linked to dopamine release and previous research has shown that increasing dopamine levels is related to reduced Pavlovian bias (34). Therefore, we hypothesized that Pavlovian bias would not only impact cessation success but also be altered by cessation. Particularly, we predicted that Pavlovian bias would increase during nicotine abstinence, reflecting a bidirectional relationship between drug cessation and Pavlovian processes.

In addition to addressing these gaps, we aimed to validate and extend previous findings on the neural underpinnings of Pavlovian-instrumental conflict. Specifically, we focused on key brain regions previously implicated in this process—the striatum, vmPFC, and ACC—to determine their involvement in different aspects of the decision-making process, including action and valence encoding. We hypothesized that greater Pavlovian bias would be associated with altered reliance on striatal and cortical activity during valuation and action selection, thereby causing maladaptive valence and action value encoding and impaired decision making.

To investigate these hypotheses, we used computational modeling and functional magnetic resonance imaging (fMRI) with the GNG task, to quantify Pavlovian bias and identify neural correlates during the smoking cessation phase, a critical period requiring smokers to learn new, adaptive instrumental behaviors, thereby potentially creating conflict with the Pavlovian system. Specifically, we used participants’ clinic participation rate as a proxy for motivation to quit and investigated its interaction with Pavlovian bias. By integrating behavioral, computational, and neuroimaging approaches, this study aims to elucidate how Pavlovian biases influence cessation outcomes and may contribute to relapse vulnerability, along with their underlying neural mechanisms.

## Methods Participants

This study was part of a project aimed at identifying markers of successful smoking cessation. We recruited 113 smokers (94 males, 15 females; ages ranging from 18 to 34 years; M=25.6, SD=3.8 years) participating in the smoking cessation clinic at the Seoul National University (SNU) Health Service Center, all provided written informed consent. The study was approved by the SNU Institutional Review Board.

Behavioral analysis included 86 participants (74 males, 12 females). Out of 113 smokers, 27 participants were excluded due to data collection issues (n=5), low concentration (sleeping inside the scanner) (n=9), excessive button presses (>90% of trials) (n=1), model convergence issues (𝑅^ > 1.1) (n=3), incomplete smoking survey during the clinic (less than 7 days out of 5 to 6 weeks) (n=9). For fMRI analyses, further exclusions were made for scanning protocol deviations (n=13 in session 1; n=12 in session 2) and mean framewise displacement (FD) > 0.2 mm (37) (n=5 in session 1; n=11 in session 2), resulting in final samples of 68 (session 1), 63 (session 2), and 57 (within-subject).

## Procedure

Participants visited the laboratory three times: eligibility screening, pre-clinic, and post-clinic. Screening was conducted online (before the first visit) and in person (first visit). All participants were daily nicotine users and met the following inclusion criteria (details in **Supplementary Table S1**): (1) no history of psychiatric illness, (2) no other substance addictions, and (3) at least 8 hours of abstinence from nicotine, alcohol, and caffeine before each visit. Those who passed our screening completed baseline surveys (see **Baseline Surveys**) at all visits. In sessions 2 and 3, participants performed the orthogonalized go/no-go (GNG) task inside the MRI scanner (see **Orthogonalized go/no-go task**). Participants received additional compensation based on task accuracy, in addition to a standard participation fee.

The smoking cessation clinic began with an education session on the dangers of smoking, benefits of quitting, and cessation strategies, followed by weekly visits for five weeks (36 days; two clinics lasted 43 days due to national holidays). Each visit included an appointment with a doctor who prescribed anti-smoking medication (Varenicline or Bupropion) or nicotine replacement therapy (NRT; nicotine patch, gum, or candy), and a counseling session based on a booklet written by the National Cancer Institute (38). Participants completed daily surveys during the clinic, using a smartphone application developed in our laboratory. The momentary smoking survey, which recorded the number of cigarettes smoked, was used to determine cessation success.

## Baseline Surveys

We measured demographics, smoking history, nicotine dependence (Fagerström Test for Nicotine Dependence; FTND (39); Kano Test for Social Nicotine Dependence; KTSND (40)), urges (Brief Questionnaire on Smoking Urges; QSU-Brief (41)), withdrawal (Minnesota Tobacco Withdrawal Scale; MTWS (42); Cigarette Withdrawal Scale-21; CWS-21 (43)), and psychological traits (Barratt Impulsiveness Scale-11; BIS-11 (44); State-Trait Anxiety Inventory Form Y; STAI-Y (45); Center for Epidemiologic Studies Depression Scale; CES-D (46); Beck Depression Inventory-II; BDI-II (47); Yale-Brown Obsessive Compulsive Symptom Checklist; Y-BOCS-SC (48)). For group models, we included KTSND, CWS-21, BIS-11, and Y-BOCS-SC as covariates to avoid multicollinearity. Detailed demographic and descriptive statistics of all baseline measures are provided in **Supplementary Table S2**.

## Assessment of smoking cessation

Abstinence was assessed through self-reports and an app-based survey, confirming no smoking during the last week. Additionally, exhaled carbon monoxide (CO) levels were measured, with levels below 5 ppm confirming abstinence (49). Individuals exceeding this threshold were excluded.

## Orthogonalized go/no-go task

The orthogonalized go/no-go (GNG) task comprises three phases per trial: cue representation, response, and outcome (**Figure 1A**). In the cue representation phase, participants saw one of four abstract fractal cues for 1000ms, each representing different trial types: press a button to gain a reward ("go to win"), press to avoid losing ("go to avoid losing"), withhold pressing to win ("no-go to win"), or withhold to avoid losing ("no-go to avoid losing"). The associations of these cues were randomized among participants.

**Figure 1.**
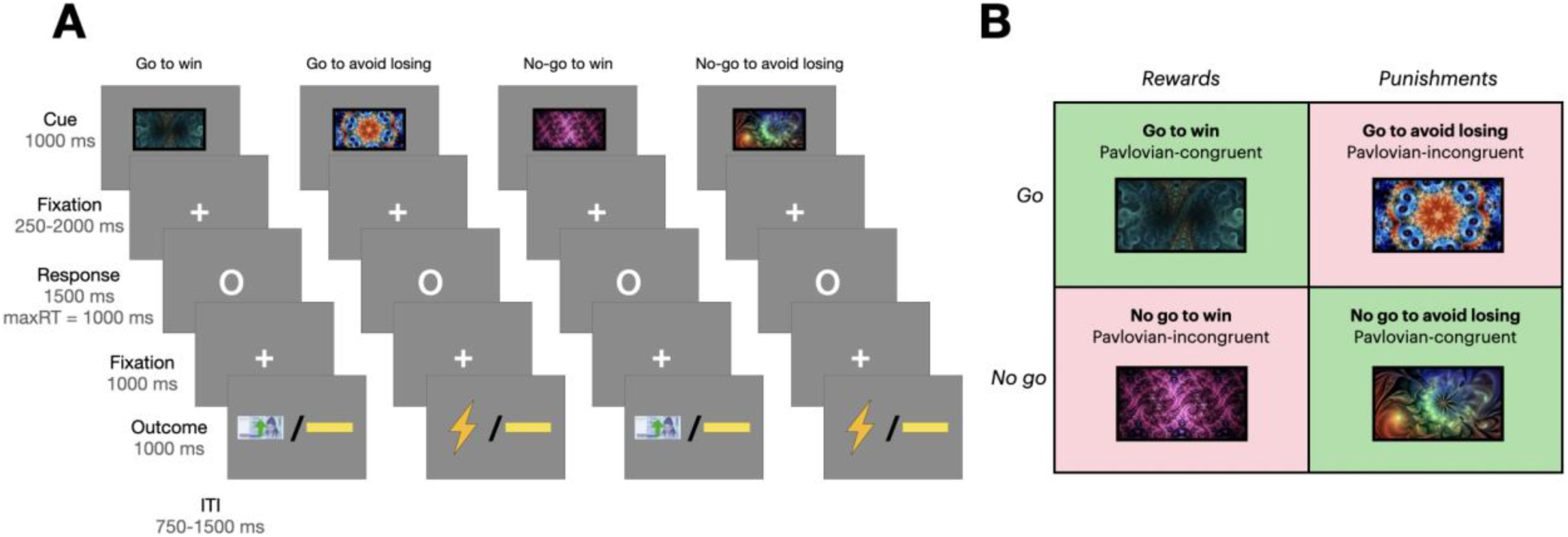
Task structure of orthogonalized go/no-go task. **(A)** Trial timeline. Each trial started with a fractal cue, followed by a jittered delay. Participants then responded to a circle by pressing a button or refraining. After another brief delay, the outcome was shown: a win of ₩ 1000 KRW (green upward arrow on a bill), an electric shock (electric image), or no outcome (yellow horizontal bar). **(B)** Four trial types. Four fractal cues indicated combinations of action (go/no-go) and outcome valence (reward/punishment).

After the cue, a white circle signaled the response phase (1000ms), during which participants had to decide whether to press a button. The outcome phase followed based on their response, showing for 1000ms a green arrow on a ₩1000 KRW bill (about 0.75 USD) for a reward, an electric shock image for a punishment, or a yellow horizontal bar for no outcome. These outcomes are probabilistic, with a 20% chance of the optimal outcome following an incorrect response.

The task included 180 randomized trials (45 per type). Punishment was administered as an electric shock to maximize the punishing effect (50,51), with shock intensity individually calibrated to a medium pain level based on participant feedback, prior to the task initiation. Actual shock was delivered with a 30% probability.

## Computational modeling

We constructed a reinforcement learning model following Guitart-Masip et al. (10), which includes distinct outcome sensitivity and Pavlovian bias for reward and punishment. This model was identified as the best-fitting model in our previous work (52). In the model, expected values 𝑄(𝑎_𝑡_, 𝑠_𝑡_) at state 𝑠 and action 𝑎 at trial 𝑡 were updated for each action on each stimulus and trial via Rescorla-Wagner rule:

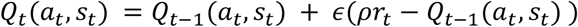

where 𝜖 represents the learning rate (53). Outcomes were entered into the equation through 𝑟_𝑡_ ∈ {1, 0, -1}, with outcome sensitivity parameters (𝜌_𝑟𝑒𝑤_, 𝜌_𝑝𝑢𝑛_) reflecting the subjective weighting of rewards and punishments.

State value 𝑉(𝑠_𝑡_) was updated similarly but independently of actions:

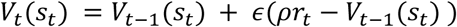

Action weights 𝑊(𝑎_𝑡_, 𝑠_𝑡_) were calculated as a combination of 𝑄(𝑎_𝑡_, 𝑠_𝑡_) and 𝑉(𝑠_𝑡_), with go bias 𝑏 and Pavlovian bias 𝜋:

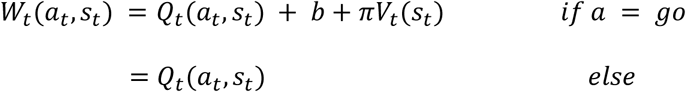

Here, 𝑏 was added to the value of go choice, and the Pavlovian bias parameter (𝜋_𝑟𝑒𝑤_ and 𝜋_𝑝𝑢𝑛_) enhanced the value of go choices in reward conditions and diminished it under punishment. Action probabilities followed a softmax function (53):

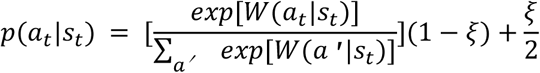

where 𝜉 represents the irreducible noise, capturing the extent to which choices were deterministically made based on the action weights.

As the study used a within-subject design across two sessions, additional delta (Δ) parameters were added to the model to capture the possible alterations between sessions. Specifically, for the post-clinic session data, each parameter included in the model was adjusted by adding a corresponding delta parameter (e.g., instead of 𝜉, 𝜉 + Δ𝜉 was used in the softmax function above), yielding 14 parameters in total.

Parameters were estimated using the hierarchical Bayesian Analysis (HBA) to improve parameter reliability through shrinkage effects (54,55), with individual estimates informed by group-level priors which were assumed to be normally distributed. The Matt trick was employed to reduce dependency among group-level parameters and facilitate the sampling process (56). Bounded parameters (e.g., irreducible noise, learning rates) were estimated in an unconstrained space and then probit-transformed to the constrained space for more efficient MCMC sampling (57). Model fitting was performed using Stan (version 2.21.8), using four independent chains (4000 samples each; 2000 warm-up samples). Convergence was verified by 𝑅^ statistics below 1.1 (**Supplementary Table S3**) and trace plot inspections (**Supplementary Figure S1**) (58).

## fMRI scans: acquisition and protocol

All scans were acquired on a Simens Tim Trio 3T scanner with a 32-channel head coil. High-resolution T1-weighted anatomical images of the whole brain resolution and functional echo-planar imaging (EPI) data were collected in three sessions. Results included in this study come from preprocessing performed using fMRIPrep 20.2.6 (58; RRID:SCR_016216), which is based on Nipype 1.7.0 (59; RRID:SCR_002502). Statistical analyses were conducted in SPM12 (Wellcome Trust Centre for Neuroimaging) under MATLAB R2018b. Full acquisition and preprocessing details are provided in **Supplementary Appendix S1**.

## fMRI scans: general linear models

We employed two distinct general linear models (GLMs) to investigate neural mechanisms underlying Pavlovian-instrumental conflicts. Specifically, GLM1 (categorical) examined action and valence effects at cue onset and their association with Pavlovian bias. GLM2 (model-based), though not discussed in detail in the following sections, further incorporated trial-wise computational regressors to assess the role of striatal and cortical regions in action and state value computation. Detailed methods and results for GLM2 are reported in the Supplementary Materials (**Appendix S2**, **Figure S5**, and **Table S8**).

For both models, participant-specific (1st-level) GLMs included six movement nuisance regressors to control for movement-related artifacts. Linear contrasts at each voxel were employed to obtain participant-specific estimates. These estimates were analyzed at the group level using a one-sample t-test, treating subjects as random effects with age and gender covariates. Differences between the quit and non-quit groups were also analyzed to assess the impact of smoking cessation on neural activity.

The categorical GLM included the following regressors: (1) cue onset of go to win trials, (2) cue onset of no-go to win trials, (3) cue onset of go to avoid losing trials, (4) cue onset of no-go to avoid losing trials, (5) target onset of go trials, (6) target onset of no-go trials, (7) outcome onset of win trials, (8) outcome onset of neutral trials, (9) outcome onset of loss trials, (10) fixation point after the outcome, and (11) wait onset, which was an inter-trial interval. Two contrasts were computed to demonstrate the action and valence-related brain activation and their association with Pavlovian bias: the main effect of action ([(1) + (3)] – [(2) + (4)]), and the main effect of valence ([(1) + (2)] – [(3) + (4)]).

## Regions of interest (ROIs)

ROIs were defined using anatomical masks from the Automatic Anatomical Labeling 3 (AAL3) atlas (61), with small volume correction (SVC) within the following ROIs.

ROIs included the striatum and substantia nigra/ventral tegmental area (SN/VTA), known to be involved in reward processing and learning (10,34). The striatum was constructed using AAL3 definitions of the bilateral caudate, putamen, olfactory bulb, and nucleus accumbens. The SN/VTA incorporated the bilateral substantia nigra and ventral tegmental area from AAL3. Additionally, cortical ROIs included the vmPFC and ACC, implicated in encoding cue valence and resolving conflicts between Pavlovian and instrumental responses (19–23).

## Results

### Behavioral performance in the GNG task

To behaviorally assess the Pavlovian bias, accuracy was computed as the percentage of correct responses per condition. Participants consistently performed better in Pavlovian congruent conditions (go to win, no-go to avoid losing) compared to incongruent conditions (no-go to win, go to avoid losing) across both sessions (**Figure 2**). Specifically, accuracy was higher in go to win than no-go to win conditions (Session 1: M=.80±.20 vs. .66±.30, t(85)=3.65, p<.001, d=0.55; Session 2: .81±.23 vs. .67±.33, t(85)=3.38, p<.001, d=0.49), and in no-go to avoid losing than go to avoid losing conditions (Session 1: .75±.13 vs. .61±.20, t(85)=5.71, p<.001, d=0.83; Session 2: .79±.15 vs. .64±.24, t(85)=5.03, p<.001, d=0.75), confirming a significant Pavlovian bias.

**Figure 2.**
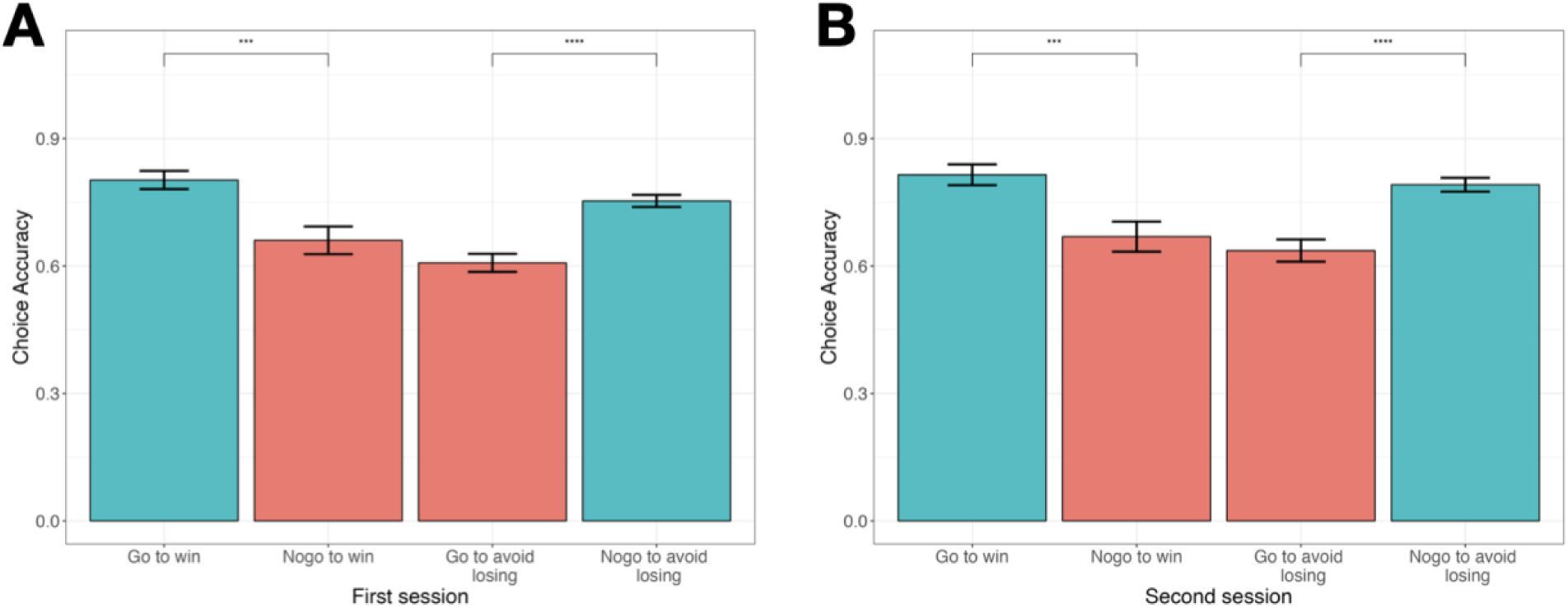
Behavioral performance in GNG task. Accuracy rates for the four conditions in the Go/No-Go (GNG) task during the first **(A)** and second **(B)** sessions. Participants showed better performance in Pavlovian congruent trials (blue) compared to incongruent trials, indicating Pavlovian bias.

### Effect of Pavlovian bias on quit attempt

First, we investigated whether Pavlovian bias measured at the baseline is directly associated with cessation success. We ran a logistic regression model predicting cessation success as a dependent variable and included Pavlovian bias (𝜋_𝑟𝑒𝑤_) estimated from computational modeling (see **Supplementary Table S4** and **Supplementary Figure S2** for detailed information of all estimated parameters), clinic participation rate, and baseline survey measures including BIS, KTSND, CWS, and YBOCS as independent variables. While clinical participation rate showed a significant main effect, increasing the log odds of cessation success (coefficient estimate=1.10, SE=0.38, p=.004), Pavlovian bias itself was not significantly associated with cessation outcome (p=.68).

Next, we further tested whether Pavlovian bias moderates the impact of motivation to quit on successful smoking cessation, using an extended logistic regression model that additionally included an interaction term between Pavlovian bias and clinic participation rate. In this model, the main effect of clinic participation remained significant (coefficient estimate=1.13, SE=0.41, p=.007), and the main effect of Pavlovian bias remained non-significant. Importantly, the interaction effect between Pavlovian bias (𝜋_𝑟𝑒𝑤_) and clinic participation rate was significant (coefficient estimate=-0.84, SE=0.39, p=.031), indicating that a higher Pavlovian bias reduced the positive effect of clinic participation on quitting (**Table 1** and **Figure 3**). This interaction effect supported our hypothesis, highlighting the crucial role of Pavlovian bias in undermining the efforts and motivation required for successful smoking cessation. Despite participants exhibiting motivation and making efforts to quit smoking, as indicated by their clinic participation rates, a higher Pavlovian bias impeded this association, rendering smoking cessation more challenging.

**Figure 3.**
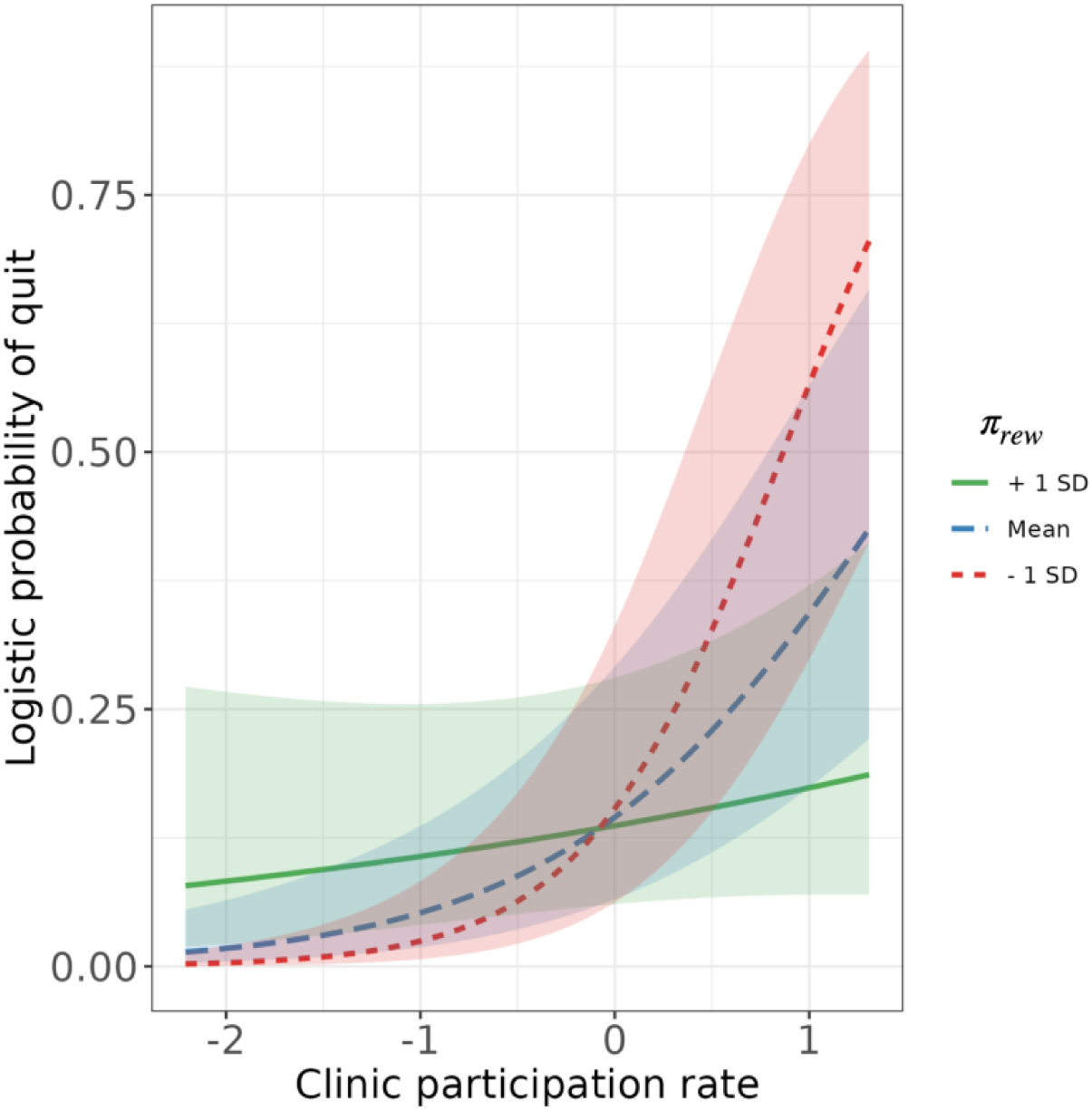
Interaction effects of participants’ Pavlovian bias and clinic participation rate Error bars indicate standard errors of the interaction effects.

**Table 1.** Logistic regression model predicting successful smoking cessation.

|  | Coefficient<br>estimate | SE | z | p-value |
| --- | --- | --- | --- | --- |
| (Intercept) | -1.775 | 0.850 | -2.087 | .037* |
| Smoking duration | 0.214 | 0.296 | 0.724 | .469 |
| Sex: male | 0.050 | 0.927 | 0.054 | .957 |
| BIS | 0.869 | 0.237 | 2.503 | .012* |
| KTSND | -0.109 | 0.325 | -0.335 | .737 |
| CWS | -1.073 | 0.435 | -2.468 | .014* |
| YBOCS | 0.366 | 0.336 | 1.089 | .276 |
| Participation rate | 1.127 | 0.414 | 2.721 | .007* |
| $\pi_{rew}$ | -0.067 | 0.337 | -0.200 | .842 |
| Participation rate x<br>$\pi_{rew}$ | -0.845 | 0.391 | -2.162 | .031* |

*Note.* BIS = Barratt Impulsiveness Scale; KTSND = Kano Test for Social Nicotine Dependence; CWS-21 = Cigarette Withdrawal Scale; YBOCS = Yale-Brown Obsessive-Compulsive Scale.

Regarding other covariates, high BIS (impulsivity) significantly increased the likelihood of quitting smoking while high CWS (cigarette withdrawal symptoms) was negatively associated with successful cessation (**Table 1**).

### Change in Pavlovian bias after smoking cessation attempt

We examined changes in Pavlovian bias from session 1 to session 2 (Δ𝜋_𝑟𝑒𝑤_ and Δ𝜋_𝑝𝑢𝑛_) to test whether Pavlovian bias increases during nicotine abstinence. Posterior distributions revealed that while quitters and non-quitters did not show a credible difference in Pavlovian bias before clinic (**Figure 4**, top), there was a credible increase in Pavlovian bias in the reward domain (Δ𝜋_𝑟𝑒𝑤_) only among quitters but not non-quitters (**Figure 4**, bottom), supporting our hypothesis that abstinence may influence Pavlovian bias and clarifying which aspect of the bias is affected.

**Figure 4.**
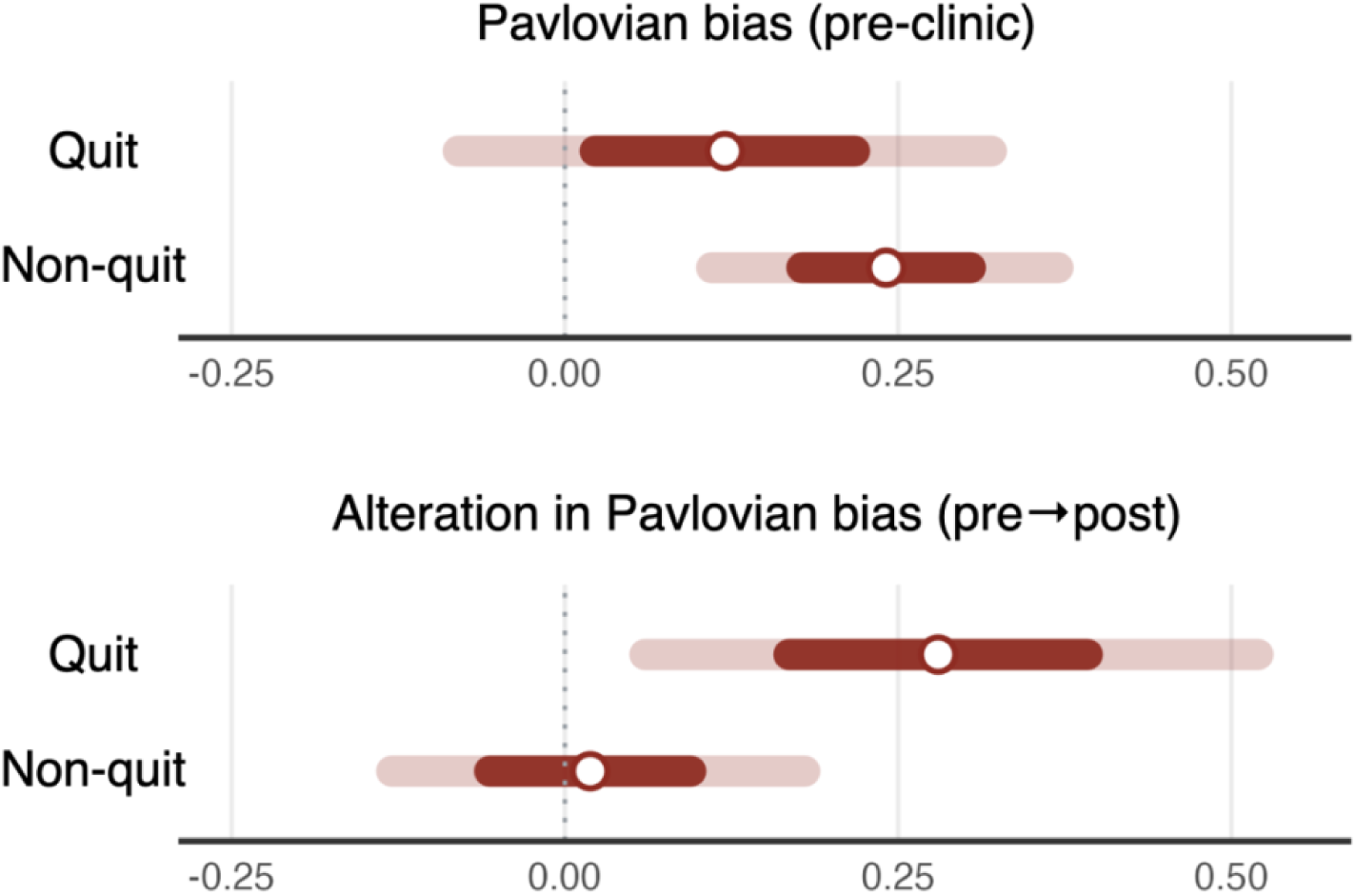
Posterior distributions of the group-level Pavlovian bias in pre-clinic session (top) and its alteration (bottom) in each group. White dots indicate medians and colored bars indicate 95% (light) and 50% (dark) highest density interval (HDI).

Furthermore, we found that non-quitters demonstrated a credible increase in reward sensitivity and irreducible noise following the cessation attempt (**Supplementary Figure S3**).

### Categorical GLM: the main effect of action and valence

Building on our behavioral results, we tested for main effects of action and valence in the cue phase and their association with Pavlovian bias (GLM1). We did not observe a significant main effect of action in our ROIs but found that no-go cues elicited greater activation than go cues in the dlPFC (MNI coordinates x=- 34, y=28, z=55, Z=4.10, p<.05, whole-brain cluster-level FWE in the first session, **Supplementary figure S4**). This activity was not significantly correlated with Pavlovian bias parameters, suggesting that the dlPFC response to action cues may not be strongly influenced by individual differences in Pavlovian bias.

In contrast, valence processing in the vmPFC was significantly modulated by Pavlovian bias. Reward-associated cues elicited greater activation in the vmPFC than punishment-associated cues (MNI coordinates x=-8, y=58, z=2, Z=5.25, p<.05, whole-brain cluster-level FWE in the first session, **Figure 5A**), consistent with previous findings (21). Importantly, individuals with higher Pavlovian bias in the reward domain (𝜋_𝑟𝑒𝑤_) exhibited reduced differentiation between reward and punishment cues in this region (r=-0.35, p=.008), indicating blunted value processing among people with high Pavlovian bias (**Figure 5B**). Because the vmPFC is crucial for integrating value information to support goal-directed control, this blunted value integration may impair the ability to appropriately integrate or regulate Pavlovian biases during decision-making.

**Figure 5.**
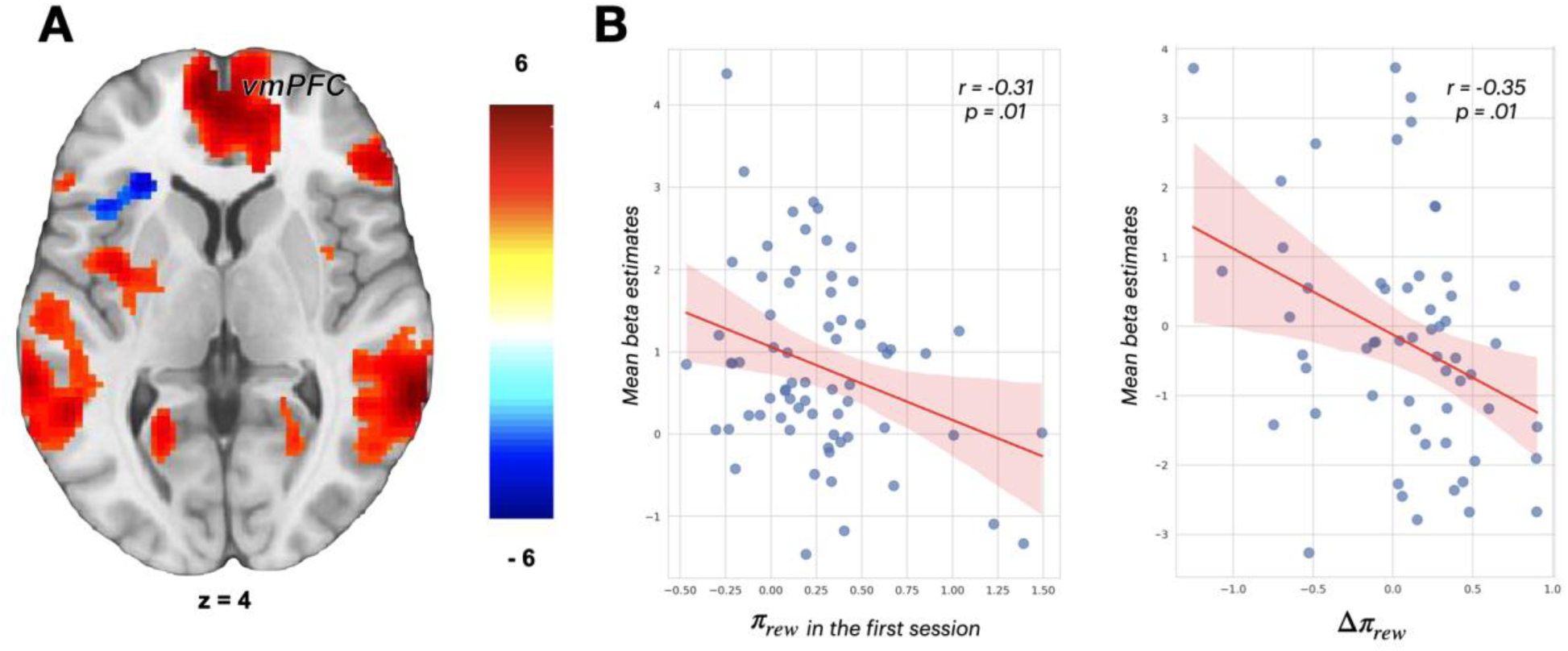
Categorical fMRI results - the main effect of valence. **(A)** Increased vmPFC activity during reward cues compared to punishment cues. Overlays are presented at a threshold of p < .001 (uncorrected). The color scale indicates t-values. **(B)** left: Correlation between 𝜋_𝑟𝑒𝑤_ in the first session and mean beta estimates extracted from the peak sphere (5mm) in the vmPFC; right: Correlation between Δ𝜋_𝑟𝑒𝑤_ and the difference in mean beta estimates extracted from the same peak sphere between sessions (ses2 – ses1). Blue dots represent individual mean beta estimates and Pavlovian bias, with the red line as the regression line. Error bars indicate standard errors.

## Discussion

This study investigated the role of Pavlovian bias in smoking cessation using the orthogonalized go/no-go task and explored underlying neural mechanisms with fMRI. Behavioral and computational modeling analyses revealed that Pavlovian bias negatively influenced the relationship between the motivation to quit and actual cessation success: higher bias reduced the benefit of treatment participation, indicating that individuals with higher Pavlovian bias are likely to face greater challenges in quitting smoking despite strong motivation and effort.

This moderating effect may help reconcile mixed findings in the literature (27,30,33,62), by clarifying that Pavlovian tendencies may not uniformly predict maladaptive behavior, but instead exert their influence particularly when strong instrumental demands conflict with automatically elicited cue-driven responses. The cessation phase provides a clear example of such conflict, as smokers must override automatic cue-driven responses to adopt new adaptive behaviors. Our findings demonstrate that Pavlovian bias can disrupt cessation efforts by moderating the relationship between motivation and successful quitting, consistent with evidence that it modulates motivational impact of conditioned cues, rather than functioning as a simple, direct driver of addiction (63). Specifically, by using clinic participation rates as indicators of motivation, we showed that Pavlovian bias toward rewards serves as a significant risk factor for cessation failure and relapse, contributing to its clinical implications.

Longitudinal analyses further revealed that successful quitters exhibited an increase in Pavlovian bias after cessation, supporting our hypothesis that nicotine abstinence heightens bias. This change may be driven by the dopamine system, which is strongly implicated in both addiction and Pavlovian bias. Previous research has demonstrated that abstinence from substances like nicotine reduces dopamine activity and increases withdrawal symptoms (36,64–66). Given that previous research has shown a link between lower dopamine levels and increased Pavlovian bias (34), it is plausible that nicotine abstinence heightens Pavlovian bias by disrupting dopamine signaling. This suggests a bidirectional relationship, where Pavlovian bias not only influences cessation success but may also be altered by the cessation process itself, particularly through the neurochemical changes induced by nicotine withdrawal.

fMRI results revealed that valence encoding in the vmPFC was significantly modulated by Pavlovian bias, with individuals exhibiting stronger Pavlovian bias in the reward domain showing reduced activation when processing the reward cues. This suggests that heightened Pavlovian tendencies may interfere with value-based processing in smokers, leading to greater reliance on automatic response patterns. This aligns with prior findings in individuals with alcohol use disorder, where reduced vmPFC activity was linked to impaired regulation of automatic appetitive responses (67), suggesting that Pavlovian bias alters valence encoding in smokers and potentially shifts decision-making toward rigid, automatic responses. Future research clarifying the role of these neural features in smoking cessation and addiction would provide insights into the mechanisms underlying relapse vulnerability and potential intervention targets.

While prior studies have emphasized the role of subcortical structures such as the striatum and midbrain in encoding Pavlovian-instrumental interactions, our findings suggest that these effects may be more condition-dependent than previously assumed. Specifically, we observed striatal activation in response to Pavlovian-incongruent cues during the first session (**Supplementary Table S9** *and* **Supplementary Figure S6**), but this effect was not associated with the individualized Pavlovian bias parameter and was not replicated in the second session. Furthermore, although we identified striatal RPE signals—initially to examine whether the basic component of reinforcement learning is blunted— these signals were only modulated by smoking cessation status, with stronger post-cessation signal reduction in individuals who successfully quit smoking (**Supplementary Table S10** and **Supplementary Figure S7**). Taken together, these results suggest that striatal involvement may emerge under specific motivational or contextual states—such as acute conflict or abstinence—rather than as a stable correlate of Pavlovian bias per se.

In conclusion, this study identifies Pavlovian bias as a significant risk factor that can hinder smoking cessation, with fMRI results showing that higher bias was associated with reduced vmPFC valence encoding. These findings underscore the need to interpret Pavlovian bias within specific motivational and contextual states. Furthermore, our findings suggest a potential cycle of smoking cessation and relapse, where nicotine abstinence increases Pavlovian bias, making sustained cessation more difficult and contributing to relapse. Integrating these neurocognitive insights into clinical practices may improve cessation outcomes and reduce relapse rates, providing a more comprehensive framework for understanding and treating addiction.

## Acknowledgments

Funding

This work was supported by the National Research Foundation of Korea (NRF) (RS-2024-00420674, RS-2025-00516410, RS-2018-NR031737) and the National Institute on Drug Abuse of the National Institutes of Health (Grant 5R01DA058038) to W.-Y.A..

## Data Availability

The data underlying this article will be shared on reasonable request to the corresponding author.

