## Supplementary Materials for "The Role of Pavlovian Bias in Smoking Cessation Success and Neurocognitive Alterations During Nicotine Withdrawal"

**Supplementary Appendix S1. fMRI acquisition protocol and preprocessing details**

Data acquisition was done on the same scanner (Simens Tim Trio 3 Tesla) using a 32-channel head coil across all participants. For spatial localization and normalization, a high resolution T1-weighted anatomical scan of the whole brain resolution was acquired for each subject (TR = 2300ms, TE = 2.36ms, FOV = 256mm, 1mm × 1mm × 1mm). Functional data was acquired using echo-planar imaging (EPI) in three scanning sessions containing 64 slices (TR = 1500ms, TE = 30ms, FOV = 256mm, 2.3mm × 2.3mm × 2.3mm). Each run consisted of 280-volume runs and lasted about 7.5 minutes.

Results included in this manuscript come from preprocessing performed using fMRIPrep 20.2.6 (58; RRID:SCR_016216), which is based on Nipype 1.7.0 (61); RRID:SCR_002502). After using fMRIPrep, images were smoothed using a 3D Gaussian kernel (8mm FWHM) for accounting for anatomical differences between participants. BOLD-signal image analysis was then performed using SPM12 [http://www.fil.ion.ucl.ac.uk/spm/] running on MATLAB v9.5.0.1067069 (R2018b).

**Anatomical data preprocessing**

A total of 2 T1-weighted (T1w) images were found within the input BIDS dataset. All of them were corrected for intensity non-uniformity (INU) with N4BiasFieldCorrection (68), distributed with ANTs 2.3.3 (61; RRID:SCR_004757). The T1w-reference was then skull-stripped with a *Nipype* implementation of the antsBrainExtraction.sh workflow (from ANTs), using OASIS30ANTs as target template. Brain tissue segmentation of cerebrospinal fluid (CSF), white-matter (WM) and gray-matter (GM) was performed on the brain-extracted T1w using fast (62; FSL 5.0.9, RRID:SCR_002823). A T1w-reference map was computed after registration of 2 T1w images (after INU-correction) using mri_-robust_template (63; FreeSurfer 6.0.1). Brain surfaces were reconstructed using recon-all (64; FreeSurfer 6.0.1, RRID:SCR_001847), and the brain mask estimated previously was refined with a custom variation of the method to reconcile ANTs-derived and FreeSurfer-derived segmentations of the cortical gray-matter of Mindboggle (65; RRID:SCR_002438). Volume-based spatial normalization to one standard space (MNI152NLin2009cAsym) was performed through nonlinear registration with antsRegistration (ANTs 2.3.3), using brain-extracted versions of both T1w reference and the T1w template. The following template was selected for spatial normalization: *ICBM 152 Nonlinear Asymmetrical template version 2009c* [(68), RRID:SCR_008796; TemplateFlow ID: MNI152NLin2009cAsym],

**Functional data preprocessing**

For each of the 8 BOLD runs found per subject (across all tasks and sessions), the following preprocessing was performed. First, a reference volume and its skull-stripped version were generated using a custom methodology of *fMRIPrep*. Susceptibility distortion correction (SDC) was omitted. The BOLD reference was then co-registered to the T1w reference using bbregister (FreeSurfer) which implements boundary-based registration (75). Co-registration was configured with six degrees of freedom. Head-motion parameters with respect to the BOLD reference (transformation matrices, and six corresponding rotation and translation parameters) are estimated before any spatiotemporal filtering using mcflirt (FSL 5.0.9, 69). BOLD runs were slice-time corrected to 0.706s (0.5 of slice acquisition range 0s-1.41s) using 3dTshift from AFNI 20160207 ((71), RRID:SCR_005927). The BOLD time-series (including slice-timing correction when applied) were resampled onto their original, native space by applying the transforms to correct for head-motion. These resampled BOLD time-series will be referred to as *preprocessed BOLD in original space*, or just *preprocessed BOLD*. The BOLD time-series were resampled into standard space, generating a *preprocessed BOLD run in MNI152NLin2009cAsym space*. First, a reference volume and its skull-stripped version were generated using a custom methodology of *fMRIPrep*. Several confounding time-series were calculated based on the *preprocessed BOLD*: framewise displacement (FD), DVARS and three region-wise global signals. FD was computed using two formulations following Power (absolute sum of relative motions, (78)) and Jenkinson (relative root mean square displacement between anes, (76)). FD and DVARS are calculated for each functional run, both using their implementations in *Nipype* (following the definitions by (78)). The three global signals are extracted within the CSF, the WM, and the whole-brain masks. Additionally, a set of physiological regressors were extracted to allow for component-based noise correction (*CompCor*, (79)). Principal components are estimated after high-pass filtering the *preprocessed BOLD* time-series (using a discrete cosine filter with 128s cut-o) for the two *CompCor* variants: temporal (tCompCor) and anatomical (aCompCor). tCompCor components are then calculated from the top 2% variable voxels within the brain mask. For aCompCor, three probabilistic masks (CSF, WM and combined CSF+WM) are generated in anatomical space. The implementation diers from that of (79) in that instead of eroding the masks by 2 pixels on BOLD space, the aCompCor masks are subtracted a mask of pixels that likely contain a volume fraction of GM. This mask is obtained by dilating a GM mask extracted from the FreeSurfer’s *aseg* segmentation, and it ensures components are not extracted from voxels containing a minimal fraction of GM. Finally, these masks are resampled into BOLD space and binarized by thresholding at 0.99 (as in the original implementation). Components are also calculated separately within the WM and CSF masks. For each CompCor decomposition, the *k* components with the largest singular values are retained, such that the retained components’ time series are sucient to explain 50 percent of variance across the nuisance mask (CSF, WM, combined, or temporal). The remaining components are dropped from consideration. The head-motion estimates calculated in the correction step were also placed within the corresponding confounds file. The confound time series derived from head motion estimates and global signals were expanded with the inclusion of temporal derivatives and quadratic terms for each (80). Frames that exceeded a threshold of 0.5 mm FD or 1.5 standardised DVARS were annotated as motion outliers. All resamplings can be performed with *a single interpolation step* by composing all the pertinent transformations (i.e. head-motion transform matrices, susceptibility distortion correction when available, and co-registrations to anatomical and output spaces). Gridded (volumetric) resamplings were performed using antsApplyTransforms (ANTs), configured with Lanczos interpolation to minimize the smoothing eects of other kernels (81). Non-gridded (surface) resamplings were performed using mri_vol2surf (FreeSurfer). First, a reference volume and its skull-stripped version were generated using a custom methodology of *fMRIPrep*. Susceptibility distortion correction (SDC) was omitted. The BOLD reference was then co-registered to the T1w reference using bbregister (FreeSurfer) which implements boundary-based registration (82). Co-registration was configured with six degrees of freedom. Head-motion parameters with respect to the BOLD reference (transformation matrices, and six corresponding rotation and translation parameters) are estimated before any spatiotemporal filtering using mcflirt (FSL 5.0.9, (76)). BOLD runs were slice-time corrected to 0.705s (0.5 of slice acquisition range 0s-1.41s) using 3dTshift from AFNI 20160207 ((77), RRID:SCR_005927). The BOLD time-series (including slice-timing correction when applied) were resampled onto their original, native space by applying the transforms to correct for head-motion. These resampled BOLD time-series will be referred to as *preprocessed BOLD in original space*, or just *preprocessed BOLD*. The BOLD time-series were resampled into standard space, generating a *preprocessed BOLD run in MNI152NLin2009cAsym space*. First, a reference volume and its skull-stripped version were generated using a custom methodology of *fMRIPrep*. Several confounding time-series were calculated based on the *preprocessed BOLD*: framewise displacement (FD), DVARS and three region-wise global signals. FD was computed using two formulations following Power (absolute sum of relative motions, (78)) and Jenkinson (relative root mean square displacement between anes, (76)). FD and DVARS are calculated for each functional run, both using their implementations in *Nipype* (following the definitions by (78)). The three global signals are extracted within the CSF, the WM, and the whole-brain masks. Additionally, a set of physiological regressors were extracted to allow for component-based noise correction (*CompCor*, (79)). Principal components are estimated after high-pass filtering the *preprocessed BOLD* time-series (using a discrete cosine filter with 128s cut-o) for the two *CompCor* variants: temporal (tCompCor) and anatomical (aCompCor). tCompCor components are then calculated from the top 2% variable voxels within the brain mask. For aCompCor, three probabilistic masks (CSF, WM and combined CSF+WM) are generated in anatomical space. The implementation diers from that of (79) in that instead of eroding the masks by 2 pixels on BOLD space, the aCompCor masks are subtracted a mask of pixels that likely contain a volume fraction of GM. This mask is obtained by dilating a GM mask extracted from the FreeSurfer’s *aseg* segmentation, and it ensures components are not extracted from voxels containing a minimal fraction of GM. Finally, these masks are resampled into BOLD space and binarized by thresholding at 0.99 (as in the original implementation). Components are also calculated separately within the WM and CSF masks. For each CompCor decomposition, the *k* components with the largest singular values are retained, such that the retained components’ time series are sucient to explain 50 percent of variance across the nuisance mask (CSF, WM, combined, or temporal). The remaining components are dropped from consideration. The head-motion estimates calculated in the correction step were also placed within the corresponding confounds file. The confound time series derived from head motion estimates and global signals were expanded with the inclusion of temporal derivatives and quadratic terms for each (83). Frames that exceeded a threshold of 0.5 mm FD or 1.5 standardised DVARS were annotated as motion outliers. All resamplings can be performed with *a single interpolation step* by composing all the pertinent transformations (i.e. head motion transform matrices, susceptibility distortion correction when available, and co-registrations to anatomical and output spaces). Gridded (volumetric) resamplings were performed using antsApplyTransforms (ANTs), configured with Lanczos interpolation to minimize the smoothing eects of other kernels (81). Non-gridded (surface) resamplings were performed using mri_vol2surf (FreeSurfer).

Many internal operations of *fMRIPrep* use *Nilearn* 0.6.2 ((78), RRID:SCR_001362), mostly within the functional processing workflow. For more details of the pipeline, see the section corresponding to workflows in *fMRIPrep*’s documentation.

**Copyright Waiver**

The above boilerplate text was automatically generated by fMRIPrep with the express intention that users should copy and paste this text into their manuscripts *unchanged*. It is released under the CC0 license.

**Supplementary Appendix S2. Model-based GLM regressors**

We additionally constructed a model-based GLM (GLM2) that included trial-wise parametric modulators to probe neural encoding of action value (Qgo, Qnogo) and state value (V(t)). Specifically, GLM2 included the following regressors: (1) cue onset of trials, (2) parametrically modulated by the expected value of action (Qgo), (3) parametrically modulated by the expected value of inaction (Qnogo), (4) target onset of go trials, (5) target onset of no-go trials, (6) outcome onset, (7) parametrically modulated by the raw outcome, (8) parametrically modulated by the state value (V (t)) of the cue stimulus, (9) fixation point after the outcome, and (10) wait onset, which was the inter-trial interval. To address brain regions which are implicated in action value encoding, we compared the main effect of Qgo and Qnogo ((2) – (3)).

**Table S1. Participant screening criteria and verification methods**

| **Criterion** | **Inclusion Rule** | **Reference/Tool** |
| --- | --- | --- |
| Nicotine use | ≥ 5 cigarettes/day or ≥ 5 e-cigarette sessions/day (≥15 puffs or ≥10 mins/session) | (79) |
| Psychiatry history | No current or past psychiatric/mental illness; no psychiatric medication use | SCID-CV (86) |
| Other addictions | No addiction to other substances (e.g., alcohol, drugs) | SCID-SV (86) |
| Recent drug use | No recent use of other substances | WHPM One-Step Drug of Abuse Test Dip Card |
| Pre-experiment abstinence | No nicotine, alcohol, or caffeine for 8 hours prior to experiment | Micro Smokerlyzer™ (CO < 12 ppm); AL8800 ALCOSCAN |

**Table S2. Summary of demographic information and psychometric and smoking-related measures**

|  | **Total** | | | **Group** | | | | | | **t** | **p** |
| --- | --- | --- | --- | --- | --- | --- | --- | --- | --- | --- | --- |
|  | **(N = 86)** | | | **Quit (N = 22)** | | | **Non-quit (N = 64)** | | |  |  |
|  | **%** | **Mean** | **SD** | **%** | **Mean** | **SD** | **%** | **Mean** | **SD** |  |  |
| Sex |  |  |  |  |  |  |  |  |  |  | 1 |
| Female | 13.95 |  |  | 9.09 |  |  | 15.63 |  |  |  |  |
| Male | 86.05 |  |  | 90.91 |  |  | 84.37 |  |  |  |  |
| Age |  | 25.69 | 3.84 |  | 25.73 | 3.81 |  | 25.67 | 3.89 | 0.06 | .954 |
| Daily smoking |  |  |  |  |  |  |  |  |  |  |  |
| 5-10 | 61.63 |  |  | 72.73 |  |  | 57.81 |  |  |  |  |
| 11-20 | 36.05 |  |  | 27.27 |  |  | 39.06 |  |  |  |  |
| >20 | 2.32 |  |  | 0 |  |  | 3.13 |  |  |  |  |
| Smoking duration |  | 5.21 | 4.27 |  | 5.45 | 4.24 |  | 5.13 | 4.31 | 0.31 | .757 |
| Clinic participation |  | 0.63 | 0.28 |  | 0.80 | 0.21 |  | 0.57 | 0.28 | 3.40 | .001 |
| BIS |  | 63.66 | 9.27 |  | 67.35 | 10.68 |  | 62.39 | 8.46 | 2.21 | .030 |
| STAI-S |  | 40.14 | 6.82 |  | 38.86 | 4.69 |  | 40.58 | 7.39 | -1.26 | .213 |
| STAI-T |  | 39.53 | 8.02 |  | 39.50 | 7.45 |  | 39.55 | 8.26 | -0.02 | .981 |
| CES |  | 12.07 | 6.95 |  | 11.27 | 5.35 |  | 12.34 | 7.44 | -0.62 | .536 |
| KBDI |  | 8.30 | 6.35 |  | 6.94 | 4.64 |  | 8.77 | 6.81 | -1.16 | .248 |
| YBOCS |  | 3.90 | 3.87 |  | 4.18 | 4.27 |  | 3.80 | 3.76 | 0.40 | .690 |
| MNWS |  | 8.98 | 6.46 |  | 7.05 | 5.58 |  | 9.64 | 6.64 | -1.64 | .104 |
| CWS |  | 42.38 | 13.64 |  | 40.14 | 9.12 |  | 43.16 | 14.86 | -1.12 | .266 |
| CDS |  | 35.32 | 7.52 |  | 34.76 | 7.13 |  | 35.52 | 7.70 | -0.41 | .686 |
| KTSND |  | 17.74 | 3.62 |  | 17.36 | 4.39 |  | 17.88 | 3.34 | -0.57 | .570 |
| QSU |  | 33.59 | 12.51 |  | 33.14 | 9.88 |  | 33.75 | 13.36 | -0.20 | .844 |
| FTND |  | 2.36 | 1.61 |  | 2.05 | 1.50 |  | 2.47 | 1.64 | -1.07 | .290 |

*Note.* A Fisher’s exact test was performed for the group difference in ‘Sex’. A Mann-Whitney test was performed for the group difference in ‘Daily smoking’. A t-test was performed for the remaining variables.

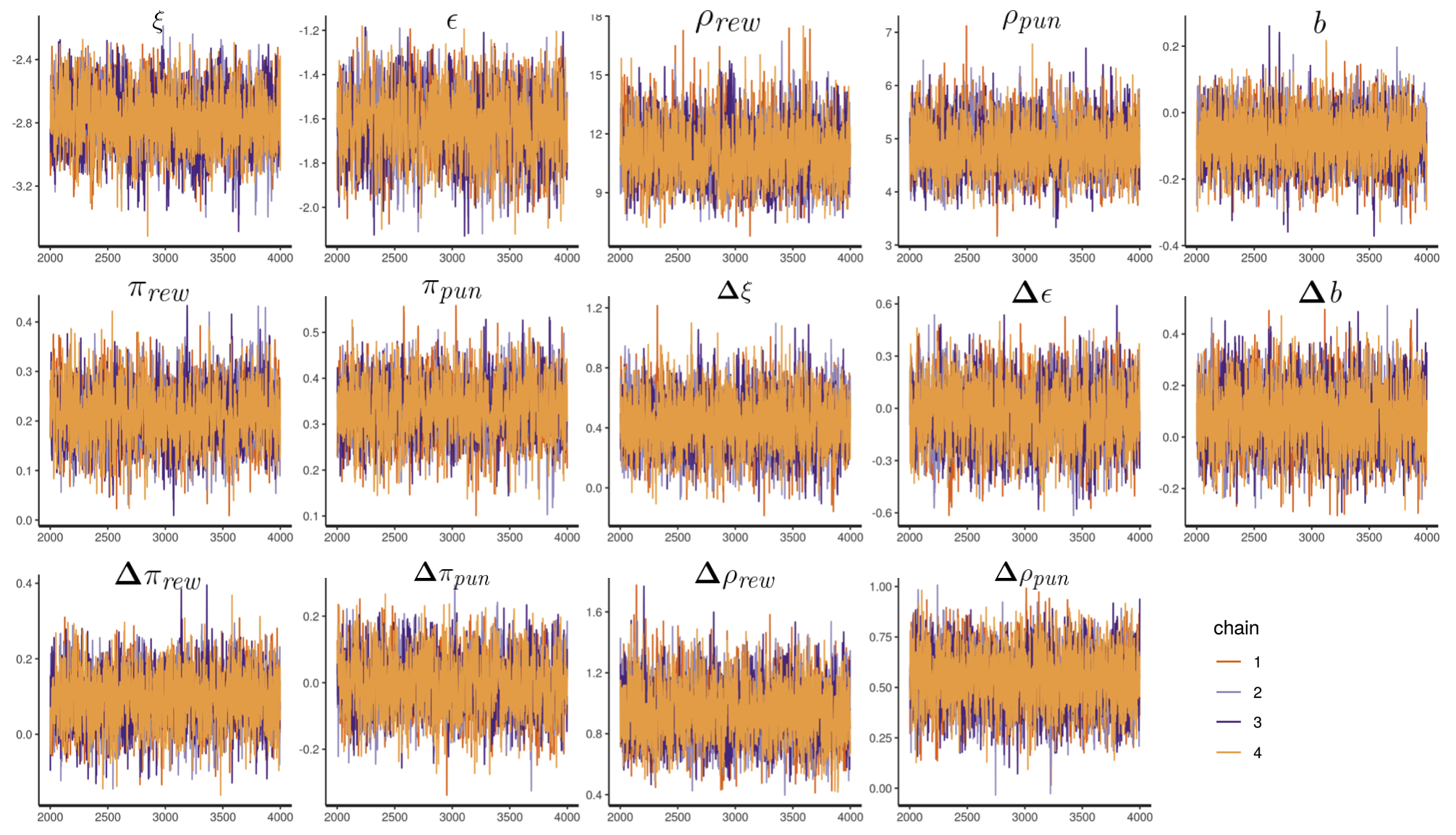

**Figure S1. Traceplots of group parameters for the model.** The traceplots show that the MCMC samples were well mixed and converged. Note that the plots excluded burn-in samples.

**Table S3. Statistics of posterior distributions of group parameters for the model**

| **Parameter** | **Mean** | **SD** | **2.5%** | **97.5%** | **R_hat** |
| --- | --- | --- | --- | --- | --- |
| $\xi$ (irreducible noise) | -2.76e+00 | 0.17 | -3.10 | -2.43 | 1.01 |
| $\epsilon$ (learning rate) | -1.63e+00 | 0.14 | -1.91 | -1.35 | 1.00 |
| $b$ (go bias) | -8.02e-02 | 0.07 | -0.23 | 0.06 | 1.00 |
| $\rho_{rew}$ (reward sensitivity) | 1.11e+01 | 1.39 | 8.68 | 14.20 | 1.00 |
| $\rho_{pun}$ (punishment sensitivity) | 4.85e+00 | 0.46 | 4.02 | 5.81 | 1.00 |
| $\pi_{rew}$ (Pavlovian bias in reward domain) | 2.15e-01 | 0.06 | 0.11 | 0.32 | 1.00 |
| $\pi_{pun}$ (Pavlovian bias in punishment domain | 3.33e-01 | 0.06 | 0.22 | 0.44 | 1.00 |
| $\Delta\xi$ (Change in irreducible noise) | 4.24e-01 | 0.17 | -0.37 | 0.32 | 1.00 |
| $\Delta\epsilon$ (Change in learning rate) | -2.66e-02 | 0.17 | -0.37 | 0.32 | 1.00 |
| $\Delta b$ (Change in go bias) | 7.80e-02 | 0.12 | -0.14 | 0.32 | 1.00 |
| ${\Delta\rho}_{rew}$ (Change in reward sensitivity) | 9.41e-01 | 0.18 | 0.60 | 1.30 | 1.00 |
| ${\Delta\rho}_{pun}$ (Change in punishment sensitivity) | 5.34e-01 | 0.14 | 0.27 | 0.81 | 1.00 |
| $\Delta\pi_{rew}$ (Change in Pavlovian bias in reward domain) | 8.98e-02 | 0.07 | -0.04 | 0.22 | 1.00 |
| $\Delta\pi_{pun}$ (Change in Pavlovian bias in punishment domain) | -1.19e-02 | 0.08 | -0.16 | 0.14 | 1.00 |

**Table S4. Statistics of posterior distributions of group parameters in the quitters**

| **Parameter** | **Mean** | **SD** | **2.5%** | **97.5%** | **R_hat** |
| --- | --- | --- | --- | --- | --- |
| $\xi$ (irreducible noise) | -2.60 | 0.20 | -3.01 | -2.23 | 1.00 |
| $\epsilon$ (learning rate) | -1.55 | -0.21 | -1.95 | -1.14 | 1.00 |
| $b$ (go bias) | -0.22 | 0.13 | -0.48 | 0.04 | 1.00 |
| $\rho_{rew}$ (reward sensitivity) | 14.31 | 2.19 | 10.73 | 19.21 | 1.00 |
| $\rho_{pun}$ (punishment sensitivity) | 4.81 | 0.63 | 4.37 | 6.17 | 1.00 |
| $\pi_{rew}$ (Pavlovian bias in reward domain) | 0.12 | 0.10 | -0.08 | 0.32 | 1.00 |
| $\pi_{pun}$ (Pavlovian bias in punishment domain | 0.24 | 0.11 | 0.03 | 0.46 | 1.00 |
| $\Delta\xi$ (Change in irreducible noise) | -0.11 | 0.20 | -0.61 | 0.40 | 1.00 |
| $\Delta\epsilon$ (Change in learning rate) | -0.05 | 0.23 | -0.49 | 0.39 | 1.00 |
| $\Delta b$ (Change in go bias) | -0.11 | 0.20 | -0.51 | 0.27 | 1.00 |
| ${\Delta\rho}_{rew}$ (Change in reward sensitivity) | 0.28 | 0.22 | -0.14 | 0.72 | 1.00 |
| ${\Delta\rho}_{pun}$ (Change in punishment sensitivity) | 0.53 | 0.19 | 0.17 | 0.90 | 1.00 |
| $\Delta\pi_{rew}$ (Change in Pavlovian bias in reward domain) | 0.28 | 0.12 | 0.06 | 0.52 | 1.00 |
| $\Delta\pi_{pun}$ (Change in Pavlovian bias in punishment domain) | -0.04 | 0.15 | -0.32 | 0.26 | 1.00 |

**Table S5. Statistics of posterior distributions of group parameters in the non-quitters**

| **Parameter** | **Mean** | **SD** | **2.5%** | **97.5%** | **R_hat** |
| --- | --- | --- | --- | --- | --- |
| $\xi$ (irreducible noise) | -2.72e+00 | 0.24 | -3.22 | -2.28 | 1.01 |
| $\epsilon$ (learning rate) | -1.59e+00 | 0.16 | -1.89 | -1.28 | 1.00 |
| $b$ (go bias) | -2.23e-02 | 0.09 | -0.19 | 0.15 | 1.00 |
| $\rho_{rew}$ (reward sensitivity) | 9.40e+00 | 1.41 | 6.94 | 12.38 | 1.00 |
| $\rho_{pun}$ (punishment sensitivity) | 4.71e+00 | 0.53 | 3.76 | 5.84 | 1.00 |
| $\pi_{rew}$ (Pavlovian bias in reward domain) | 2.41e-01 | 0.07 | 0.11 | 0.37 | 1.00 |
| $\pi_{pun}$ (Pavlovian bias in punishment domain | 3.63e-01 | 0.07 | 0.22 | 0.50 | 1.00 |
| $\Delta\xi$ (Change in irreducible noise) | 6.47e-01 | 0.22 | 0.23 | 1.10 | 1.00 |
| $\Delta\epsilon$ (Change in learning rate) | -5.58e-02 | 0.19 | -0.43 | 0.31 | 1.00 |
| $\Delta b$ (Change in go bias) | 1.42e-01 | 0.13 | -0.11 | 0.40 | 1.00 |
| ${\Delta\rho}_{rew}$ (Change in reward sensitivity) | 1.22e+00 | 0.19 | 0.85 | 1.62 | 1.00 |
| ${\Delta\rho}_{pun}$ (Change in punishment sensitivity) | 5.76e-01 | 0.16 | 0.27 | 0.89 | 1.00 |
| $\Delta\pi_{rew}$ (Change in Pavlovian bias in reward domain) | 1.90e-02 | 0.08 | -0.13 | 0.18 | 1.00 |
| $\Delta\pi_{pun}$ (Change in Pavlovian bias in punishment domain) | -1.07e-02 | 0.09 | -0.19 | 0.17 | 1.00 |

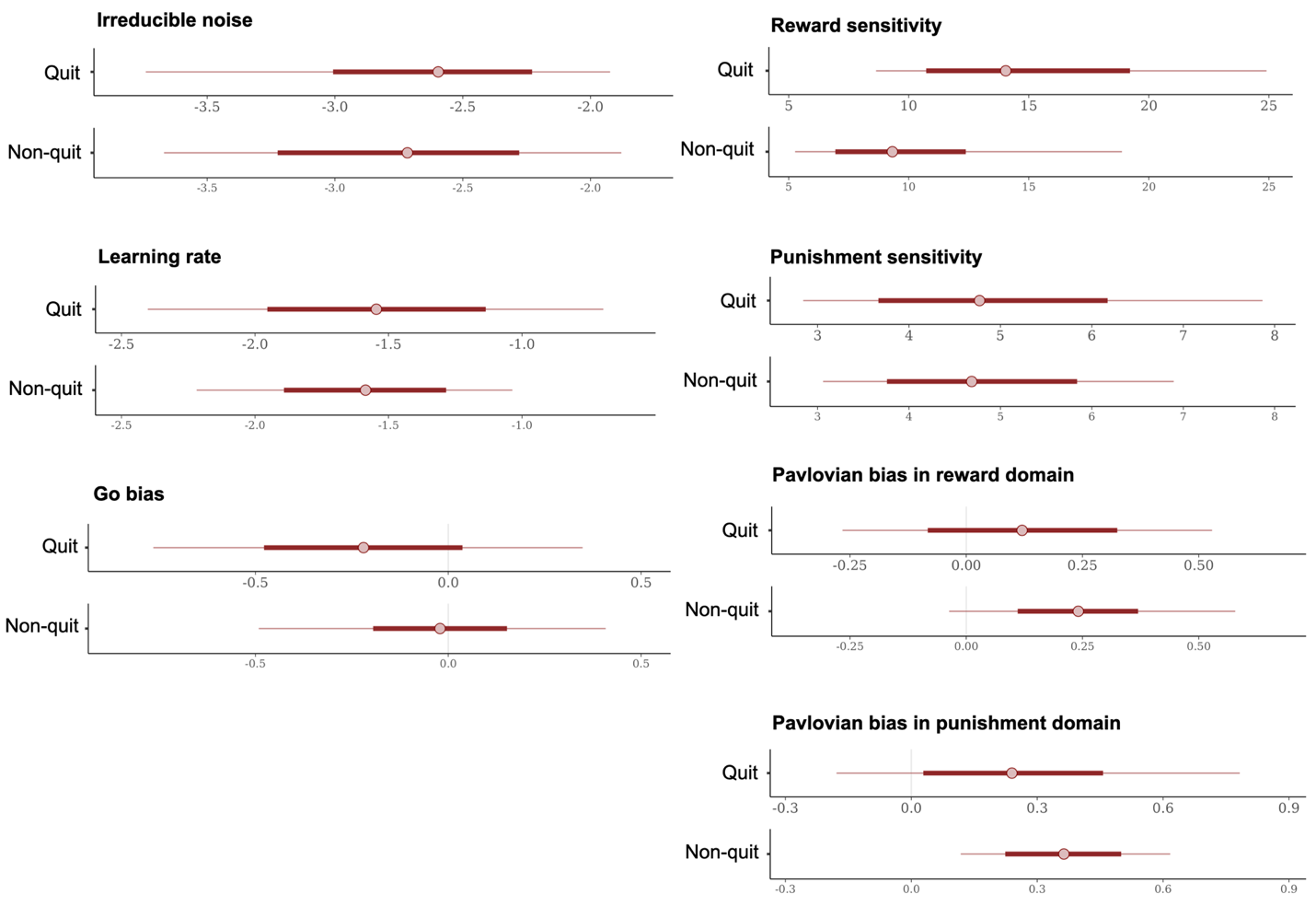

**Figure S2. Posterior distributions of the group-level parameters in each group.** Dots indicate medians and thick bars indicate 95% highest density interval (HDI).

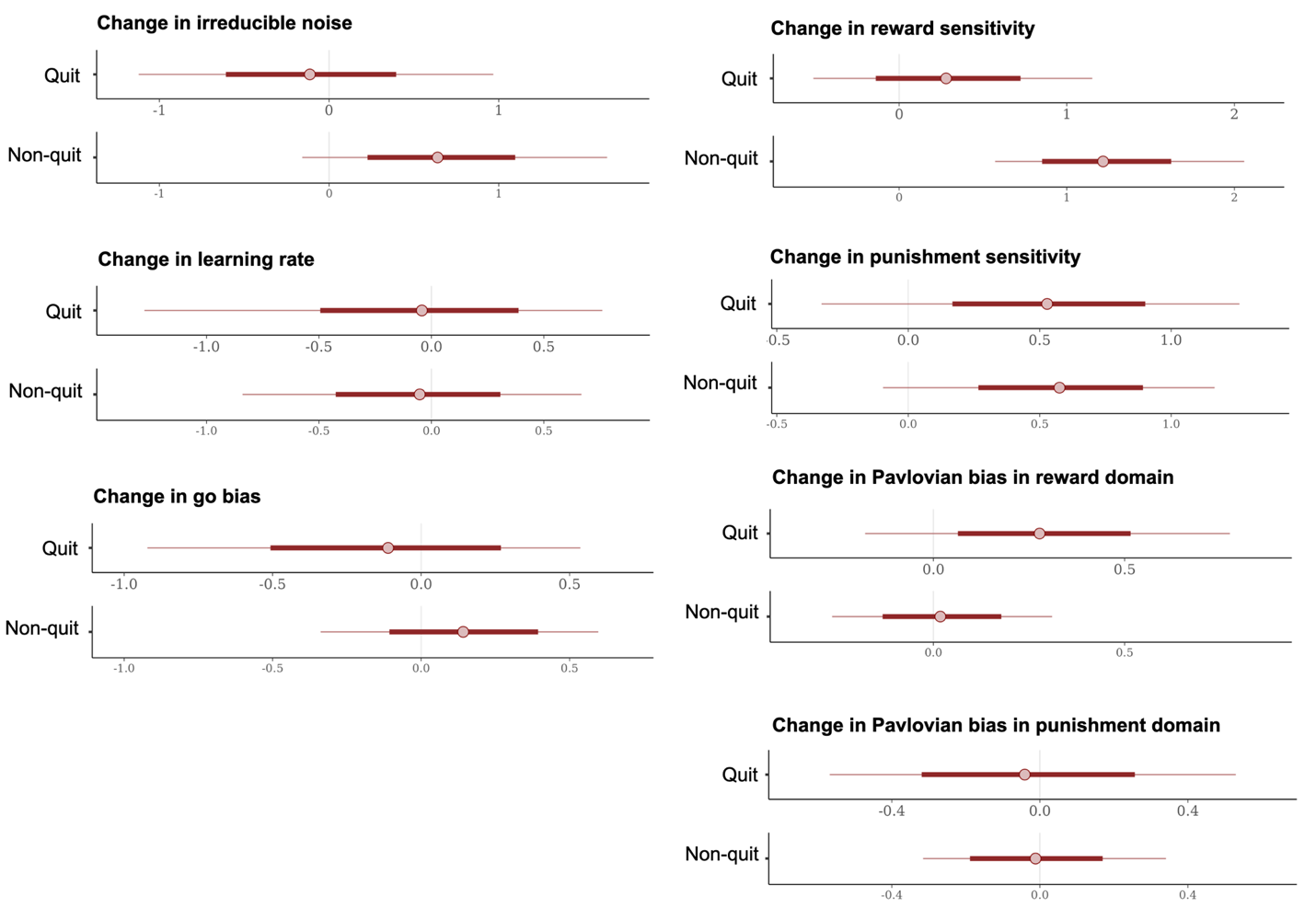

**Figure S3. Posterior distributions of the group-level delta parameters in each group.** Dots indicate medians and thick bars indicate 95% highest density interval (HDI).

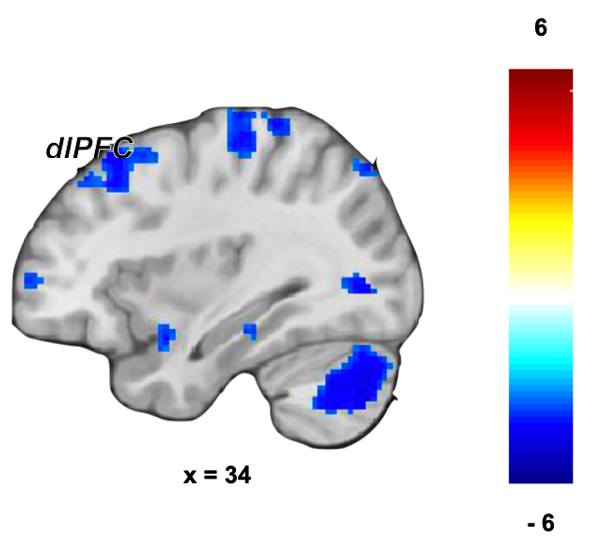

**Figure S4. Categorical GLM results – the main effect of action.** Overlays are presented at a threshold of p < .001 (uncorrected). The color scale indicates t values.

**Table S6. Whole-brain results from categorical GLM analysis (the main effect of action) in the first session**

| **Peak in region** | **Number of voxels** | **Peak voxel** | | | | **p*** |
| --- | --- | --- | --- | --- | --- | --- |
|  |  | x (mm) | y (mm) | z (mm) | Z |  |
| L cerebrum | 12439 | 15 | -21 | -16 | 5.35 | .000 |
| R inferior temporal gyrus | 453 | 50 | -28 | -19 | 4.84 | .001 |
| R anterior cingulate gyrus | 321 | 11 | 28 | 9 | 4.77 | .005 |
| L anterior cingulate gyrus | 490 | -8 | 26 | 11 | 4.72 | .001 |
| L amygdala | 336 | -15 | -2 | -19 | 4.43 | .004 |
| L middle frontal gyrus | 484 | -43 | 16 | 43 | 4.34 | .001 |
| L superior frontal gyrus | 159 | -24 | 63 | 18 | 4.19 | .078 |
| R middle frontal gyrus | 508 | 34 | 28 | 55 | 4.10 | .000 |
| R angular gyrus | 426 | 55 | -63 | 34 | 3.77 | .001 |
| R middle frontal gyrus | 196 | 45 | 47 | 11 | 3.58 | .040 |

*Whole-brain cluster-level family-wise error (FWE) for multiple comparison with a cluster-forming threshold of p < .001.

**Table S7. Whole-brain results from categorical GLM analysis (the main effect of valence) in the first session**

| **Peak in region** | **Number of voxels** | **Peak voxel** | | | | **p*** |
| --- | --- | --- | --- | --- | --- | --- |
|  |  | x (mm) | y (mm) | z (mm) | Z |  |
| L postcentral gyrus | 21256 | -41 | -35 | 57 | 6.97 | .000 |
| L inferior frontal gyrus | 326 | -48 | 30 | -12 | 5.26 | .003 |
| L superior frontal gyrus | 4499 | -8 | 58 | 2 | 5.25 | .000 |
| R postcentral gyrus | 926 | 52 | -21 | 43 | 5.24 | .000 |
| R lateral orbital gyrus | 185 | 36 | 37 | -10 | 4.59 | .039 |
| R inferior frontal gyrus | 210 | 52 | 35 | 4 | 4.50 | .025 |

*Whole-brain cluster-level family-wise error (FWE) for multiple comparison with a cluster-forming threshold of p < .001.

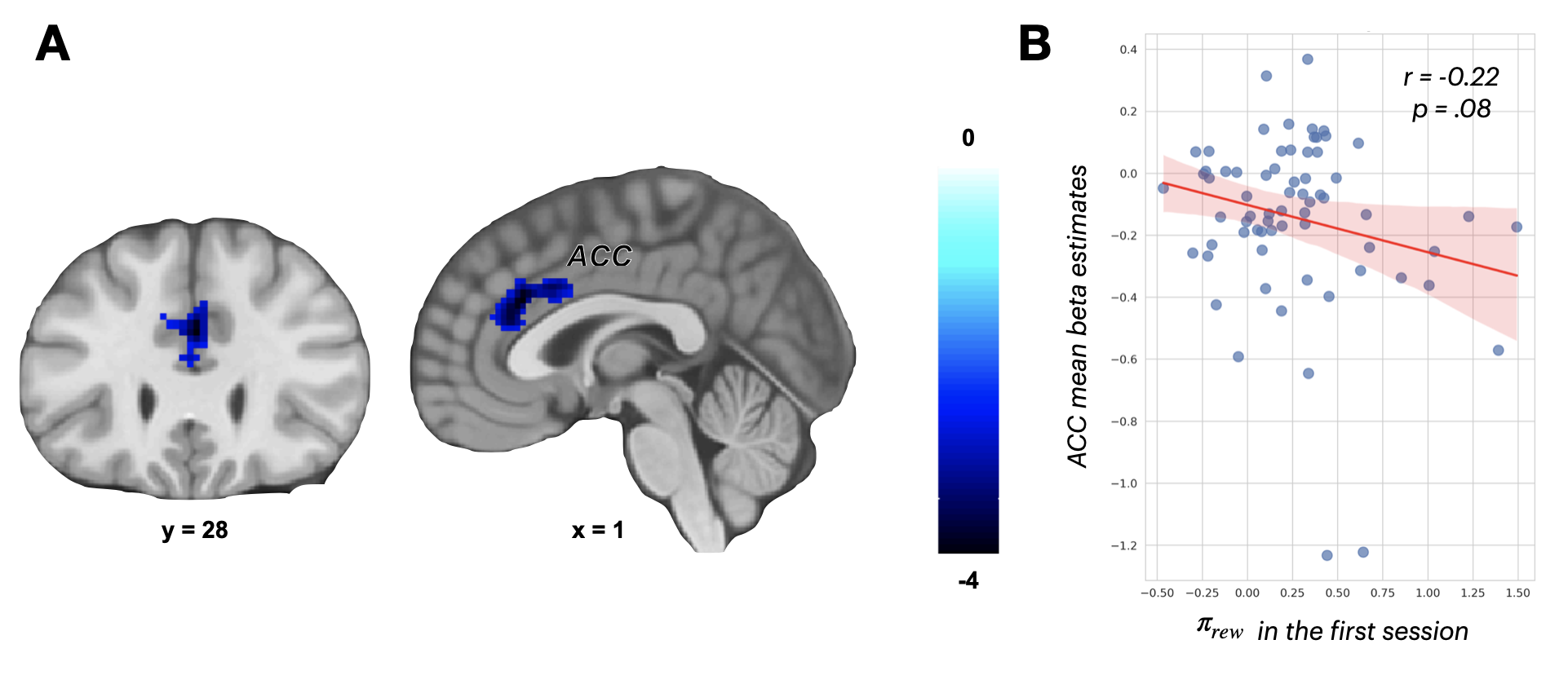

**Figure S5. Model-based fMRI results - the main effect of action value** **(A)** Negative relationship between ACC activation and action value ($Q_{go}-Q_{nogo}$) in the first session, with overlays at p < .001 (uncorrected). Color scale indicates t-values. **(B)** Correlation between mean beta estimates extracted from the peak sphere (5mm) in the ACC and Pavlovian bias ($\pi_{rew}$). Red dots represent individual mean beta estimates and Pavlovian bias, with the blue line as the regression line. Error bars indicate standard errors.

**Table S8. ROI results from model-based GLM analysis (the main effect of action value) in the first session**

| **Peak in region** | **Number of voxels** | **Peak voxel** | | | | **p*** |
| --- | --- | --- | --- | --- | --- | --- |
|  |  | x (mm) | y (mm) | z (mm) | Z |  |
| R anterior cingulate gyrus | 126 | 1 | 28 | 27 | 4.53 | .003 |

*Small-volume corrected FWE within an anatomical ACC ROI defined from aal3 atlas with a cluster-forming threshold of p < .001.

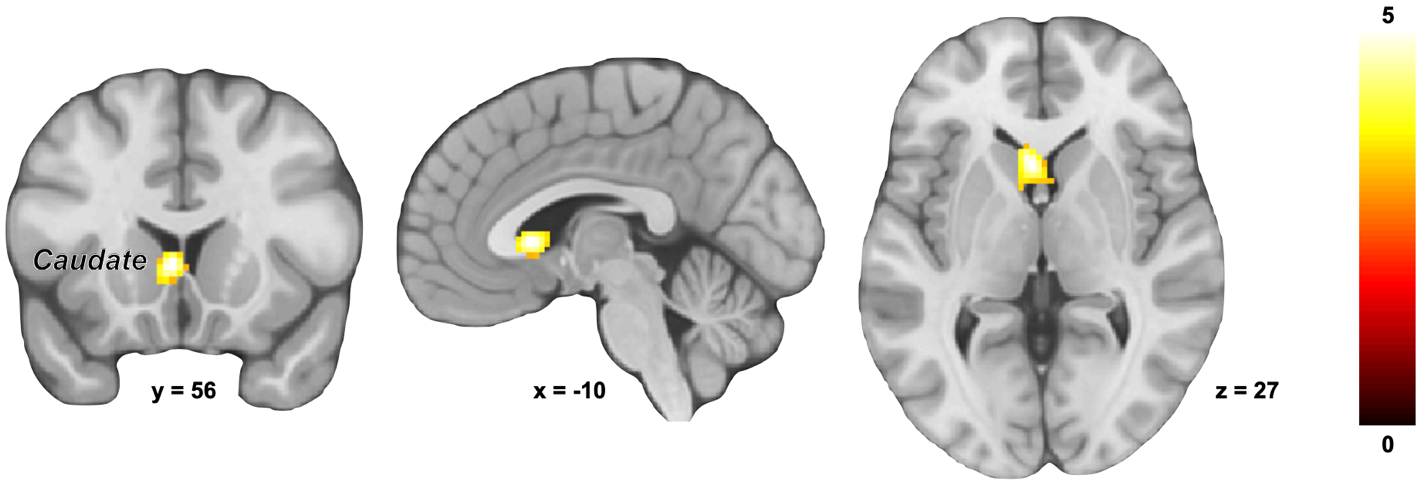

**Figure S6. Categorical GLM results – the main effect of Pavlovian bias (congruent – incongruent conditions).** Overlays are presented at a threshold of p < .001 (uncorrected). The color scale indicates t values.

**Table S9. ROI results from categorical GLM analysis (the main effect of Pavlovian bias) in the first session**

| **Peak in region** | **Number of voxels** | **Peak voxel** | | | | **p*** |
| --- | --- | --- | --- | --- | --- | --- |
|  |  | x (mm) | y (mm) | z (mm) | Z |  |
| L caudate | 78 | -3 | 16 | 2 | 4.73 | .020 |

*Small-volume corrected FWE within an anatomical striatum and SN/VTA ROI defined from aal3 atlas with a cluster-forming threshold of p < .001.

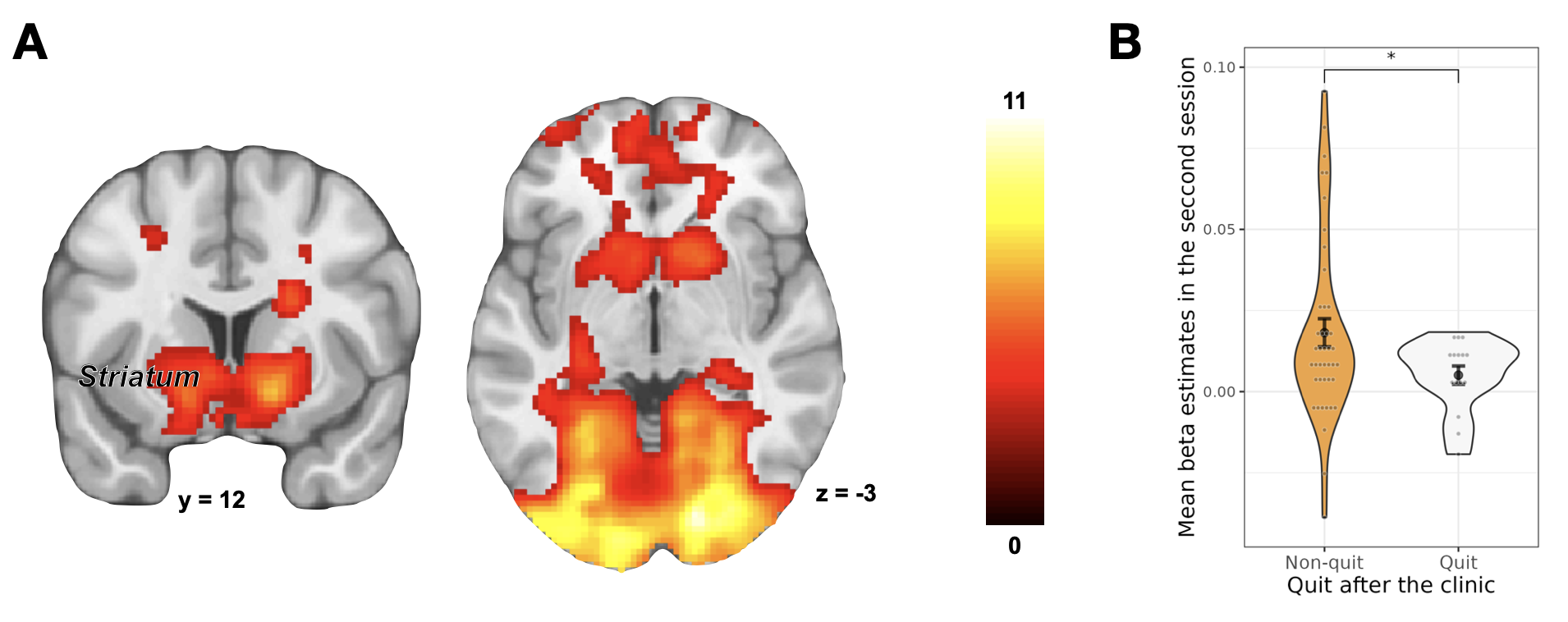

**Figure S7. Model-based GLM results – the main effect of RPE. (A)** Increased RP signaling in the striatum and **(B)** Group difference in the mean beta estimates in the post-clinic session. Overlays are presented at a threshold of p < .001 (uncorrected). The color scale indicates t values. The gray dots are mean beta estimates of individuals, the black dots are group means, and black error bars are ±SEM.

**Table S10. Whole-brain results from model-based GLM analysis (the main effect of RPE)**

| **Session** | **Peak in region** | **Number of voxels** | **Peak voxel** | | | | **p*** |
| --- | --- | --- | --- | --- | --- | --- | --- |
|  |  |  | x (mm) | y (mm) | z (mm) | Z |  |
| First session (before clinic) | R occipital fusiform gyrus | 17657 | 22 | -74 | -12 | Inf | .000 |
|  | L superior frontal gyrus | 408 | -17 | 44 | 48 | 6.79 | .000 |
|  | R putamen | 189 | 15 | 9 | -10 | 6.35 | .000 |
|  | L frontal pole | 169 | -6 | 70 | 9 | 6.08 | .000 |
|  | L inferior temporal gyrus | 167 | -62 | -35 | -21 | 5.67 | .000 |
|  | R superior frontal gyrus | 96 | 4 | 47 | -10 | 5.62 | .000 |
|  | L putamen | 108 | -15 | 14 | -12 | 5.43 | .000 |
| Second session (after clinic) | L middle frontal gyrus | 1096 | -38 | 54 | 4 | 6.13 | .000 |
|  | R cerebellum | 622 | 34 | -67 | -28 | 6.11 | .000 |
|  | L cerebellum | 700 | -41 | -72 | -26 | 6.00 | .000 |
|  | L supramarginal gyrus | 730 | -52 | -49 | 46 | 5.98 | .000 |
|  | L caudate | 166 | -82 | 2 | 11 | 5.70 | .000 |
|  | L superior frontal gyrus | 341 | -3 | 26 | 34 | 5.59 | .000 |
|  | R putamen | 54 | 18 | 12 | -12 | 5.50 | .000 |
|  | R middle frontal gyrus | 161 | 31 | 63 | 2 | 5.50 | .000 |
|  | L putamen | 65 | -20 | 16 | -10 | 5.32 | .000 |

*Whole-brain cluster-level family-wise error (FWE) for multiple comparison with a cluster-forming threshold of p < .001.
